# Improved prediction of antimicrobial resistance in *Klebsiella pneumoniae* using machine learning

**DOI:** 10.1101/2024.12.10.627815

**Authors:** Jordi Sevilla-Fortuny, Fernando González-Candelas, Neris García-González

## Abstract

*Klebsiella pneumoniae* is an important cause of healthcare-associated infections, with high levels of antimicrobial resistance (AMR) to critical antibiotics such as carbapenems and third-generation cephalosporins (3GCs). Accurate antimicrobial susceptibility detection is essential for guiding appropriate treatment. In this study, we evaluated the efficacy of machine learning (ML) models for predicting AMR phenotypes in *K. pneumoniae* particularly for antibiotics for which rule-based approaches fail. We analyzed a dataset of 5,907 *K. pneumoniae* genomes from public databases and a genomic surveillance project in Spanish hospitals. ML models were trained to predict AMR phenotypes using genomic features, and their performance was compared to ResFinder, which implements a conventional rule-based approach. Models were evaluated based on predictive accuracy across antibiotics. Additionally, we conducted a detailed analysis of the genomic features associated with AMR identified by ML to identify new putative AMR determinants. ResFinder exhibited low prediction accuracy for amikacin, fosfomycin, and piperacillin/tazobactam, whereas ML models significantly improved the prediction accuracy for these antibiotics. Moreover, we provide insights into why rule-based methods failed in these cases, specifically related to the genes *acc(6)-Ib-cr*, *fosA*, and *bla*_OXA-1_, respectively. Finally, we found possible genetic factors related to resistance for each antibiotic. Our findings underscore the value of ML models in AMR prediction based on genome information for *K. pneumoniae*, especially in challenging cases where traditional methods have low success rates. Continued evaluation and refinement of ML approaches are essential for applying these methods to enhance AMR detection in clinical and public health contexts.

**Importance:** To combat antimicrobial resistance (AMR), the rapid and accurate identification of resistance phenotypes is essential for guiding appropriate therapy. In this study, we demonstrate the significant potential of machine learning (ML) to improve AMR prediction in *Klebsiella pneumoniae* using genomic data. Our findings reveal that gold standard rule-based methods for predicting AMR from genomic data underperform for antibiotics such as amikacin, fosfomycin, and piperacillin/tazobactam. In this study, we identified the genomic determinants that mislead resistance predictions in rule-based methods providing insights that can refine existing rule-based approaches. Moreover we used ML models that improved the prediction accuracy for these antibiotics and used these models to uncover potential new AMR-associated genes that contribute to a deeper understanding of resistance mechanisms. While these findings are specific to *K. pneumoniae*, the ML approach is broadly applicable to other pathogens facing similar challenges, enabling improved AMR prediction without reliance on prior knowledge.

## Introduction

*Klebsiella pneumoniae* is a major cause of healthcare-associated infections, with increasing rates of antimicrobial resistance (AMR) particularly to carbapenems and third-generation cephalosporins (3GCs) (1, 2). The rise of resistant strains poses a critical challenge for infection control and treatment, as resistance to these essential antibiotics severely limits therapeutic options. This limitation contributes to higher mortality rates (1, 3), prolonged hospitalizations, and, consequently, increased healthcare costs (4). As treatment options are limited, rapid and accurate AMR detection is crucial to guide appropriate antimicrobial therapy and effectively manage infections caused by this pathogen (5).

The growing accessibility of whole-genome sequencing (WGS) has facilitated genomic prediction of AMR as a potential alternative to traditional Antimicrobial Susceptibility Testing (AST) (6–8). The most widely used approach, commonly referred to as rule-based prediction, relies on identifying known AMR determinants in bacterial genomes. For *Klebsiella* species, popular rule-based tools include ResFinder (9), AMRFinderPlus (10), and Kleborate (11). ResFinder have been widely used to predict resistance in *K. pneumoniae*, achieving high accuracies specially for ß-lactams. Yet, predictions for other antibiotics such as amikacin or piperacillin-tazobactam are less accurate and highly variable depending on the studied dataset (6, 12, 13). Moreover, the rule-based approach performs poorly for novel resistance mechanisms or antibiotics for which the genetic basis of resistance in *K. pneumoniae* is not completely known (14).

Recent advancements have introduced machine learning (ML) as a method to predict AMR phenotypes from WGS data without relying on pre-existing knowledge, offering a more comprehensive and unbiased prediction of resistance (15–18). ML models have been successfully applied to predict multi-drug resistance in *Mycobacterium tuberculosis* (19) and nontyphoidal *Salmonella* with accuracies greater than 0.90 (16). In *E. coli* ML models have helped to discover new β-lactamase substrate activities (20), highlighting the potential of ML to reveal new mechanisms underlying AMR (21). For *K. pneumoniae, a* few studies have implemented ML models, demonstrating improved prediction accuracy over rule-based methods for several antibiotics (15, 22, 23). Nevertheless, these studies often use datasets restricted to a few clonal groups or specific geographic regions, which may limit the generalizability of the models. Despite its advantages, ML implementation requires large, high-quality datasets containing both genomic information and the corresponding phenotypic AMR profiles, which can be a significant limitation.

In this study, we aim to apply ML models to improve the accuracy of AMR phenotype prediction in cases where rule-based strategies have a low performance. To address this, we have leveraged on the genomic and phenotypic information of a comprehensive dataset comprising both publicly available *K. pneumoniae* genomes and a genomic surveillance collection from Spanish hospitals. By applying ML models to this diverse dataset, we have assessed not only the overall improvement in prediction accuracy but have also identified genetic determinants contributing to resistance to these antibiotics.

## Results

### Genome collection

We collected a total of 4,487 *K. pneumoniae* genomes with associated MIC data from the PATRIC database to which we added 1,605 genomes from the SKPCV project. After filtering, a total of 5,907 genomes were retained for further analysis.

These genomes were mainly collected in 3 different continents, Europe (n= 2,968, 50%), Asia (n= 456, 7.7%) and America (n= 1,996, 33.7%) (Figure 1A and B); and corresponded mainly to clinical samples from humans (n= 5,507, 93%). The collection was highly diverse in terms of AMR profiles (Figure 1C) and lineages. We found 435 different STs (Figures 1D and E). However, the ten most prevalent STs included 61% of the isolates. ST307 was the most prevalent type with 1,199 genomes, followed by ST258 (n=708), ST11 (n=412), and ST15 (n=341) (Figure 1E). Some STs, such as ST258, ST512, or ST11, were enriched in carbapenemase coding genes while others, such as ST307, were enriched in extended-spectrum beta-lactamases (ESBL) genes (Figure 1D).

**Figure 1.**
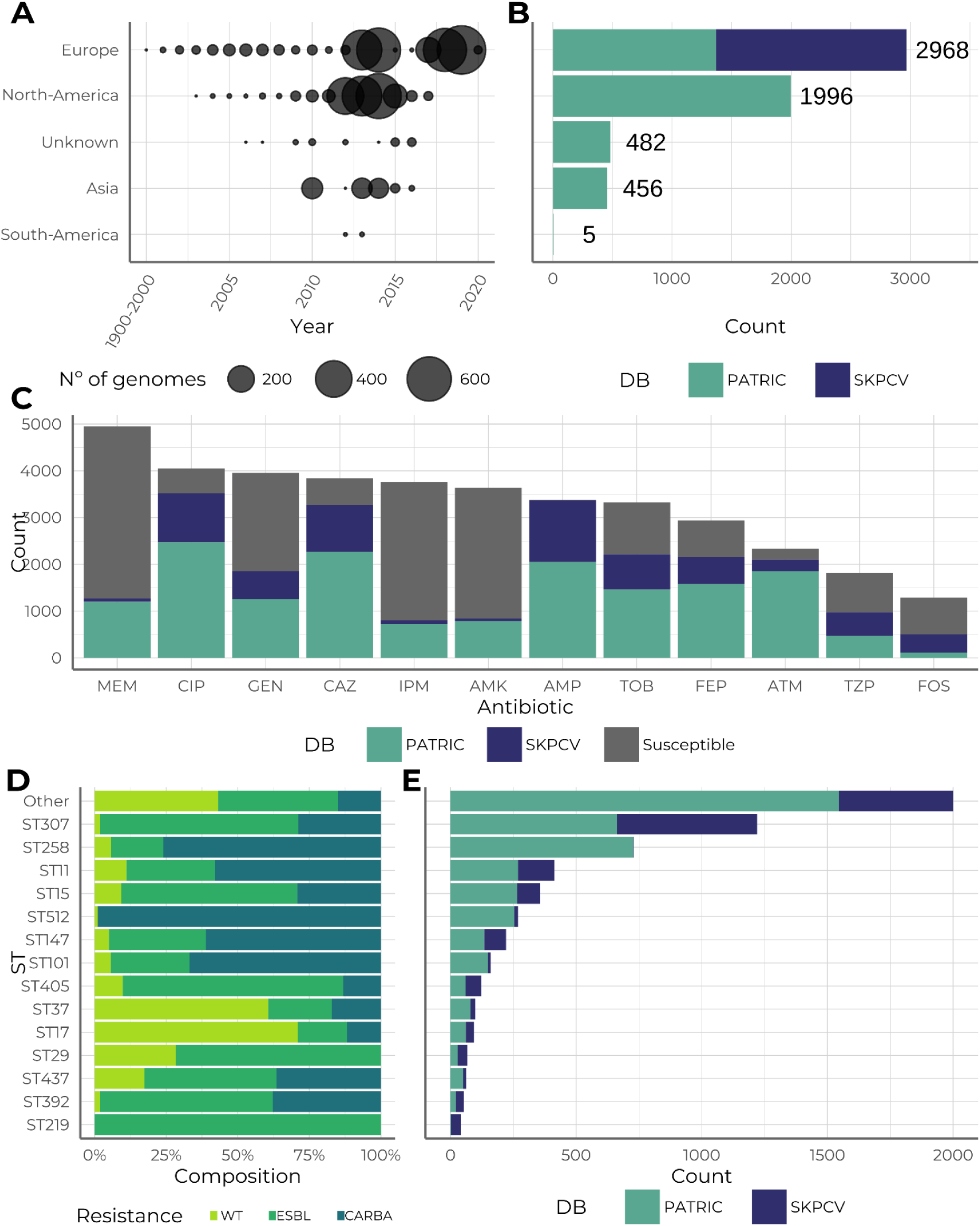
Description of the genome collection. A) Temporal and geographic distribution of the collection genomes. B) Number of genomes per geographic region and source, PATRIC or SKPCV. C) Number of tested isolates for each antibiotic. Resistant isolates are coloured by the source of the isolates, green bars represent genomes from the PATRIC database, while blue bars represent genomes from the SKPCV project. Grey represents susceptible isolates from both collections. Abbreviations: AMK = amikacin, AMP = ampicillin, ATM = aztreonam, CAZ = ceftazidime, CIP = ciprofloxacin, FEP = cefepime, FOS = fosfomycin, GEN = gentamicin, IPM = imipenem, MEM = meropenem, TOB = tobramycin, TZP = piperacillin/tazobactam. D) Distribution of major sequence types (STs) along with the presence of carbapenemase genes (CARBA), extended-spectrum β-lactamase (ESBL) genes. Wild Type (WT) indicates the absence of these genes. E) Number of genomes in each ST. Green bars represent genomes from the PATRIC database, while blue bars represent genomes from the SKPCV project.

On average, 7 antibiotics (range 0 - 12) were tested per isolate. A total of 3,383 (57.27%) isolates were multidrug-resistant (MDR) isolates (24). Resistance levels varied significantly between antibiotics, ranging from 13% to 99%. As expected, nearly all isolates were resistant to ampicillin (n=3,369, 99.7%). High resistance rates were also noted for some beta-lactams such as aztreonam (89%), cefepime (73%), and ceftazidime (85%). In contrast, carbapenems showed a lower resistance rate, with 21% for imipenem and 25% for meropenem (Figure 1C).

### Rule based prediction fails for fosfomycin, amikacin and piperacillin/tazobactam

ResFinder’s performance varied significantly across different antibiotics, with prediction accuracies ranging from 39% to 96% (Table 1). The best performances were achieved for carbapenems and aminoglycosides. Specifically, both imipenem and meropenem achieved accuracy, TPR, and TNR values exceeding 0.80. Gentamicin and tobramycin predictions also demonstrated high performance, with accuracy values of 0.91 and 0.85, respectively. In contrast, resistances to third-generation cephalosporins were poorly predicted, particularly true negative rates, which were 0.44 for ceftazidime and 0.19 for cefepime. The lowest performances in terms of accuracy were observed for the prediction of resistances to amikacin, fosfomycin, and piperacillin/tazobactam, (0.59, 0.39, and 0.57, respectively; Table 1).

**Table 1.** Prediction of the resistant phenotype using ResFinder for the 12 selected antibiotics. TPR = true positive rate, TNR = true negative rate.

| Antibiotic | TPR | TNR | Accuracy (95% CI) | F1-score |
| --- | --- | --- | --- | --- |
| Amikacin | 0.82 | 0.52 | 0.59 (0.58, 0.61) | 0.48 |
| Ampicillin | 0.96 | 0.60 | 0.96 (0.96, 0.97) | 0.98 |
| Aztreonam | 0.97 | 0.34 | 0.91 (0.90, 0.92) | 0.95 |
| Ceftazidime | 0.97 | 0.44 | 0.89 (0.88, 0.90) | 0.94 |
| Ciprofloxacin | 0.98 | 0.03 | 0.86 (0.85, 0.87) | 0.92 |
| Cefepime | 0.98 | 0.19 | 0.77 (0.76, 0.79) | 0.86 |
| Fosfomycin | 0.99 | 0.00 | 0.39 (0.36, 0.42) | 0.56 |
| Gentamicin | 0.89 | 0.93 | 0.91 (0.90, 0.92) | 0.90 |
| Imipenem | 0.94 | 0.83 | 0.85 (0.84, 0.86) | 0.73 |
| Meropenem | 0.94 | 0.80 | 0.84 (0.83, 0.85) | 0.75 |
| Tobramycin | 0.91 | 0.74 | 0.85 (0.84, 0.86) | 0.89 |
| piperacillin/tazobactam | 0.69 | 0.44 | 0.57 (0.55, 0.60) | 0.63 |

### ML improves the prediction of antimicrobial resistance

The rule-based prediction could not achieve accuracies above 75% for amikacin, fosfomycin, and piperacillin/tazobactam resistance. Therefore, we trained ML predictive models for these antibiotics to evaluate whether predictive performance could be improved.

First, we optimized the model parameters using gentamicin because this was one of the most frequently tested antibiotics in our dataset. We explored 32 different combinations of learning rate and subsampling across 10 sets of data. Since the data did not follow a normal distribution, we performed marginal Kruskal-Wallis tests. We found no significant impact of subsampling (df = 7, p = 0.08) or learning rate (df = 3, p = 0.94) on AUC performance at the 95% confidence. However, subsampling had a significant effect on AUC variance (*Levene’s test,* df = 7, p-value = 0.004). The lowest standard deviation was obtained for a subsampling level of 0.7 (sd = 0.015), which was selected for further analyses. For the learning rate, since no significant difference in variance was detected (Levene’s test, df = 3, p = 0.78), we opted for a value of 0.06, in line with previous studies (25). With the hyperparameters established, we trained predictive models for resistance to amikacin, fosfomycin, and piperacillin/tazobactam. The ML models performed significantly better than the rule-based prediction strategy for the three antibiotics based on accuracy.

Amikacin resistance was predicted using 1,125 genomes for training and 375 for testing. The model reached an AUC of 0.95 (Figure 2A). The optimal threshold to discretize the prediction was estimated to be 0.054, leading to an accuracy of 0.90 (95% CI = [0.86, 0.93]), a bAcc of 0.92, and an F1-score of 0.83. The prediction for piperacillin/tazobactam resistance was done using 1,125 genomes for training and 375 for testing. It showed an AUC of 0.86 (Figure 2A), and an optimal threshold of 0.39, resulting in an accuracy of 0.80 (95% CI = [0.76, 0.83]), bAcc of 0.80, and F1-score of 0.83. Fosfomycin prediction was done using 968 genomes for training and 321 for testing. The performance of the model achieved an AUC of 0.78 (Figure 2A). The optimal threshold was estimated to be 0.38, entailing an accuracy of 0.74 (95% CI = [0.69, 0.79]), a bAcc of 0.73, and an F1-score of 0.67.

**Figure 2.**
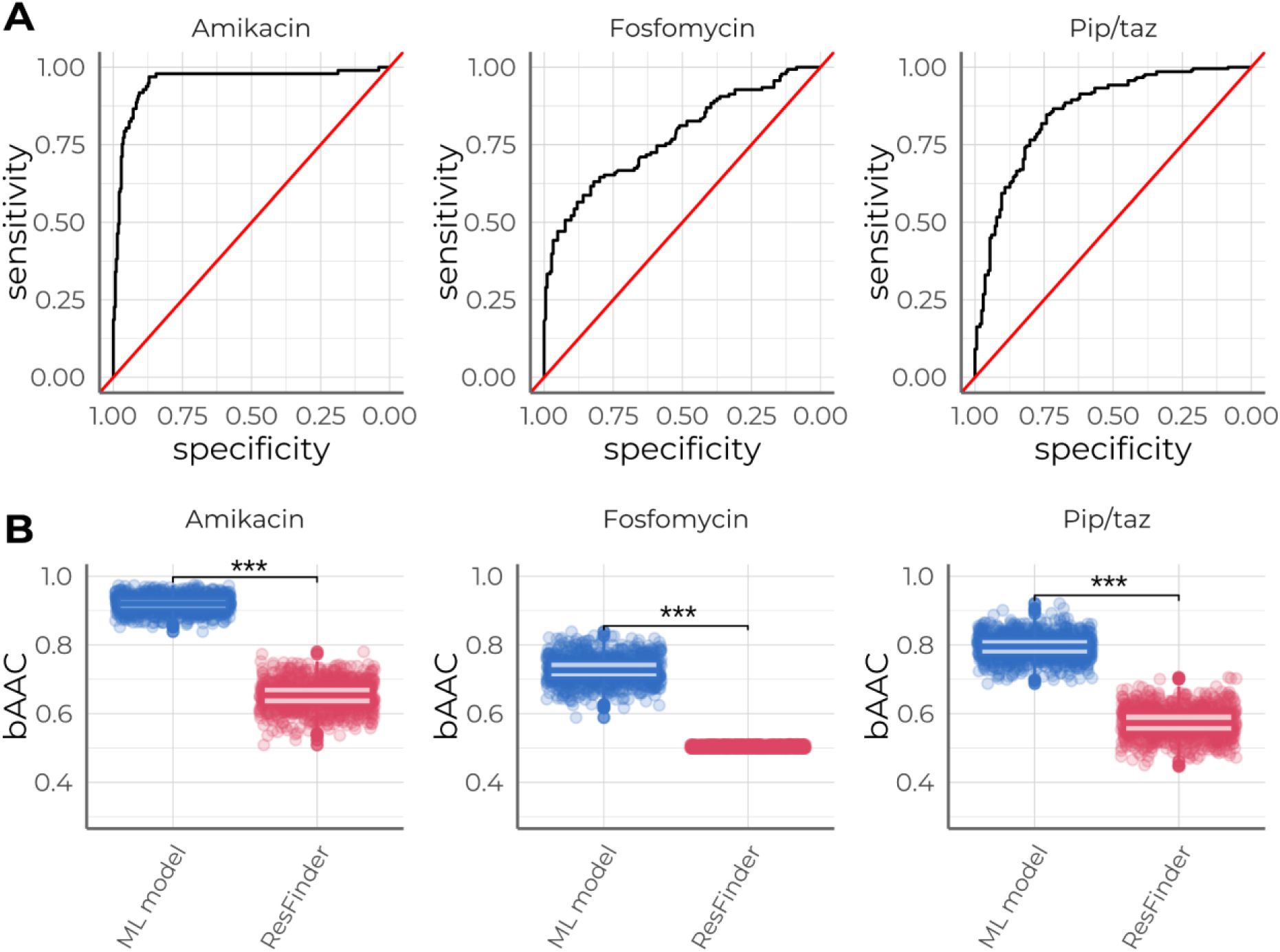
Performance for the machine learning (ML) models trained. A) ROC curves depicting the ability of ML models to predict resistance to amikacin, fosfomycin, and piperacillin/tazobactam using the testing datasets. The red line represents the ROC curve for a random classifier. B) Comparison of balanced accuracy (bAcc) for predicting resistance to amikacin, fosfomycin, and piperacillin/tazobactam between the ML models and ResFinder. Significance levels: ***p < 0.01.

For the three antibiotics, ML gave a significantly higher bAcc than ResFinder. The average bAcc for amikacin using the ML strategy was 0.92, which was significantly higher than the mean bAcc obtained with the rule-based strategy, 0.65 (Wilcoxon’s test, p < 0.001). The mean bAcc for piperacillin/tazobactam using the ML strategy was 0.80, and 0.58 using the rule-based strategy (Wilcoxon’s test, p < 0.001). For fosfomycin the ML strategy obtained a mean bAcc of 0.73 and the rule-based strategy yielded 0.50 (Wilcoxon’s test, p < 0.001) (Figure 2B; Table S1).

### Deciphering genomic determinants of antimicrobial resistance

To explore the location and genetic features that ML models found as predictors of resistance to amikacin, fosfomycin, and piperacillin/tazobactam, we identified the 10 *k*-mers with highest relative importance in each ML model (Table 2). Then, we estimated their SHAP values, which quantify the impact of a specific feature on a model’s prediction (Figure 3). These *k*-mers were used in searches to identify the genes associated with resistance. In this analysis, we also included the filtered *k*-mers identified as their duplicates in the training data. However, these duplicated k-mers were either overlapping or complementary annotations, leading to no distinct gene annotations. The results of the *k*-mer annotations are summarized in Table S2.

**Figure 3.**
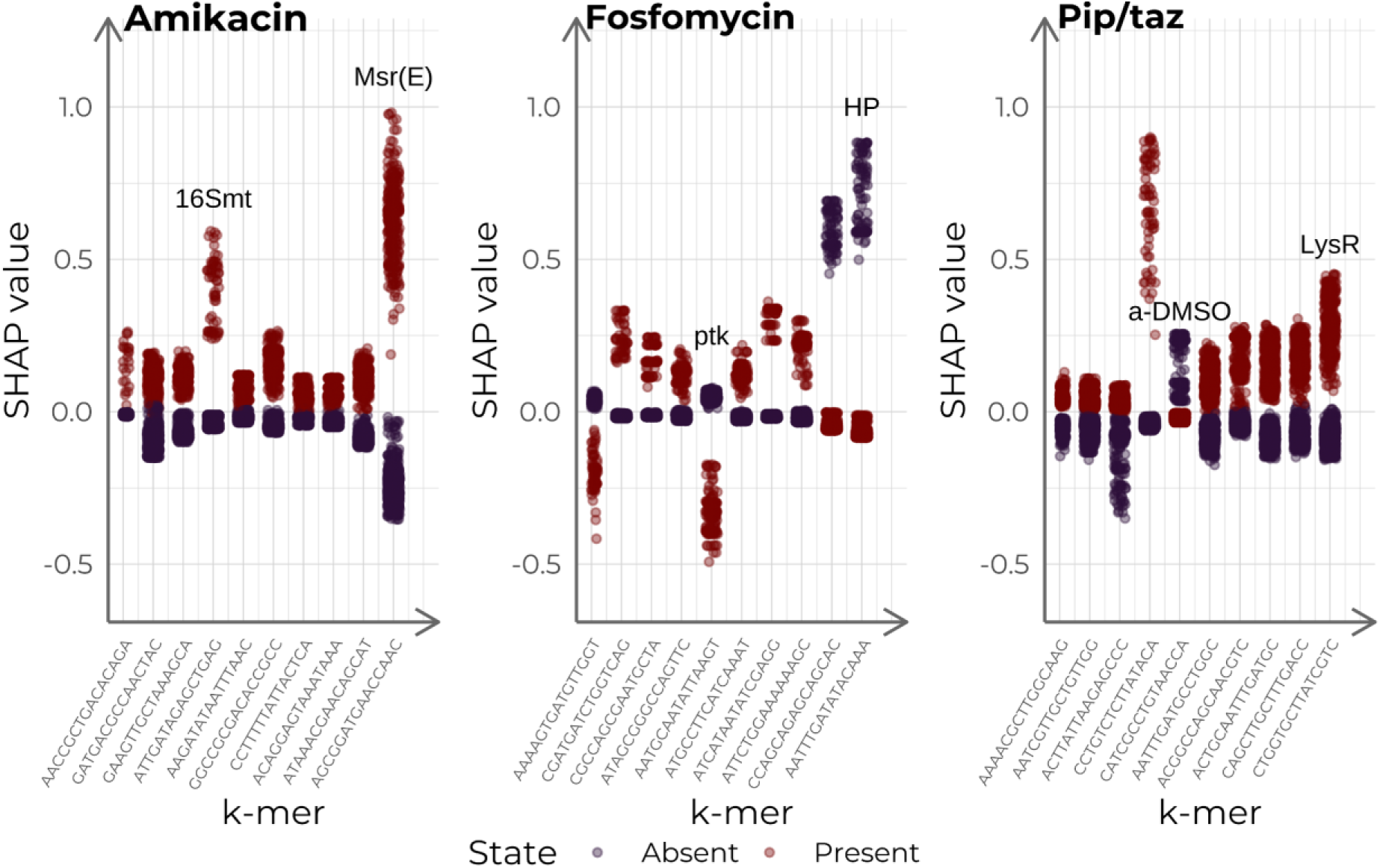
Distribution of SHAP values for the ten most important features of each model. Features in the x-axis in ascending order of importance. Dot colors represent whether it is a SHAP value for the absence (blue) or presence (red) of the feature. Function annotations for *k*-mers are shown in the plot. 16Smt = 16S rRNA (guanine(1405)-N(7))-methyltransferase, ptk = putative tyrosine-protein kinase protein, HP = hypothetical protein, a-DMSO = anaerobic dimethyl sulfoxide reductase.

**Table 2.**
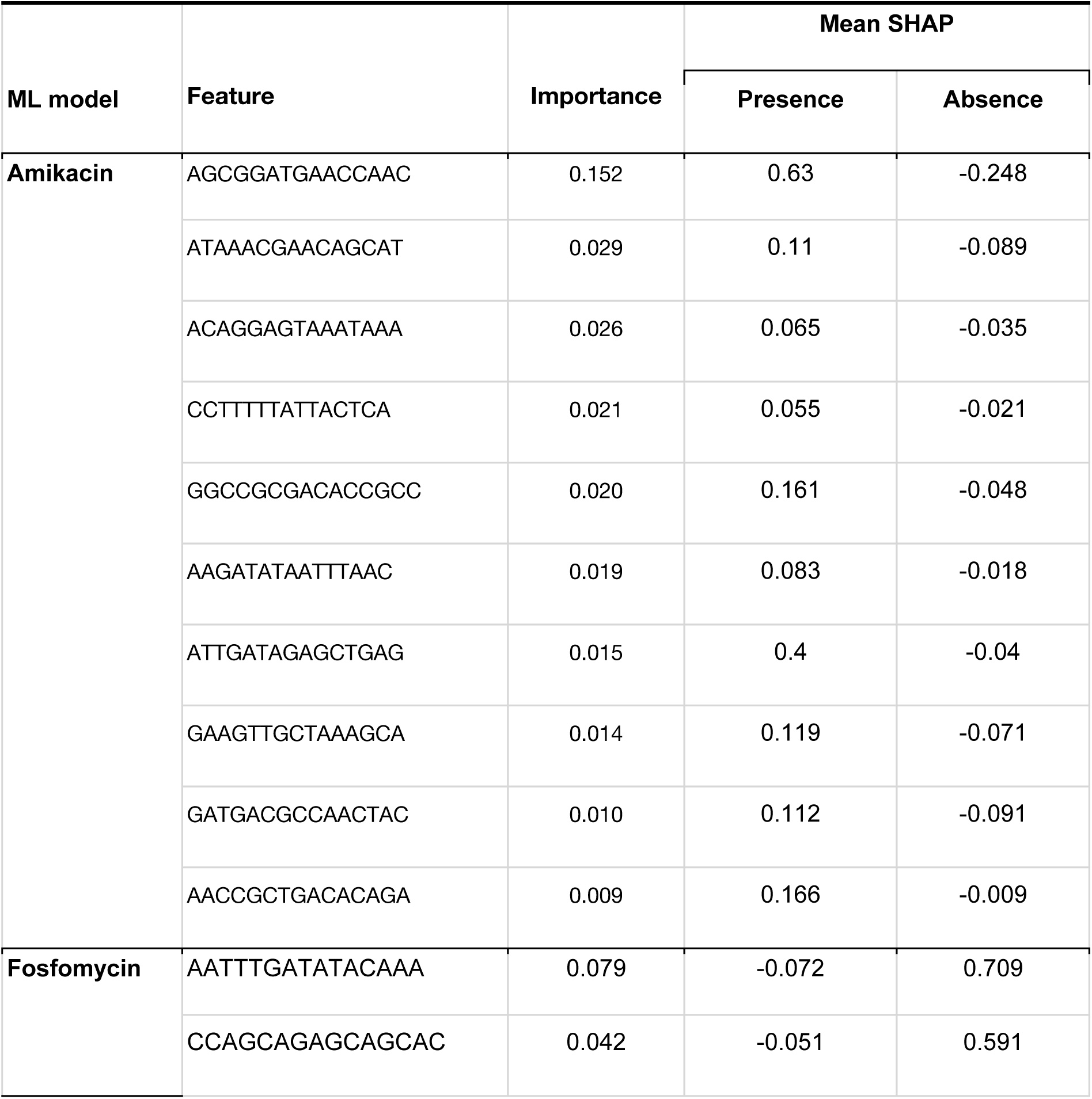
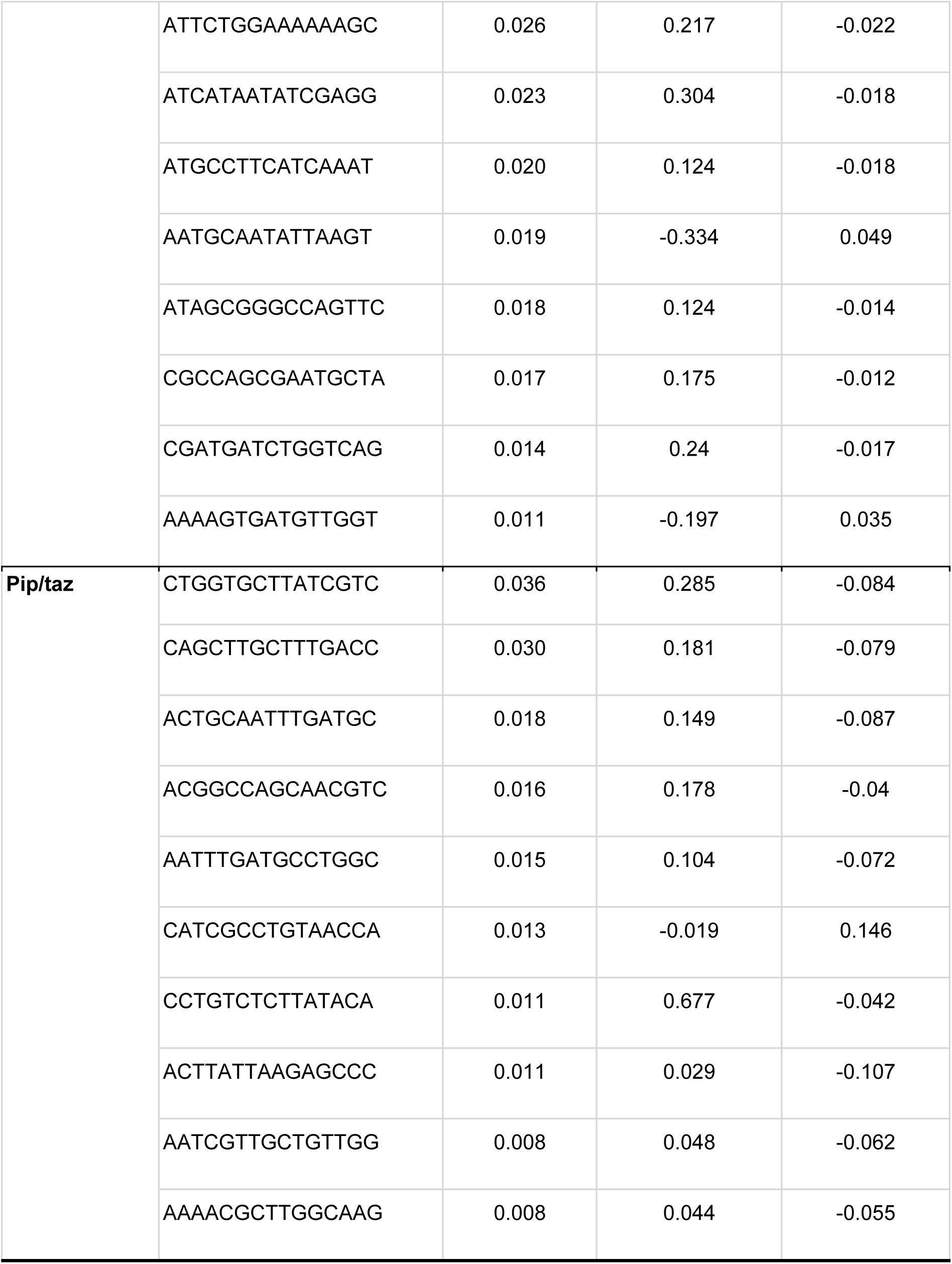
Relative importance and SHAP values for the top 10 most important features (15-mers) for the resistant phenotype prediction of the three antibiotics studied. Pip/taz = piperacillin/tazobactam.

For amikacin and fosfomycin, the most important *k*-mer showed significantly greater importance than the others, being 5.24 and 1.88 times more influential than the second-ranked *k*-mer, respectively. In contrast, for piperacillin/tazobactam, the two most important *k*-mers had similar importance (0.036 and 0.030, respectively), but a notable decrease in importance was observed starting with the third *k*-mer, which had an importance of 0.018 (Table 2).

For amikacin, ResFinder exhibited a high false positive rate (FPR = 0.48). Most of the misclassifications (n= 1,077, 72.3%) were primarily due to susceptible isolates classified as resistant due to the presence of the *AAC(6’)-Ib-cr* gene, which encodes an aminoglycoside N-acetyltransferase enzyme from the *aac(6’)* family. Removing this gene from the ruled-based prediction notably increased the accuracy (0.87) and the bAcc (0.84). However, it was still significantly lower compared to the ML model (W*ilcoxon’s test*, p < 0.001). Remarkably, none of the important *k*-mers in the ML model corresponded to the *aac(6’)* gene (Figure 3). The *k-*mer with the highest importance was annotated in two regions. The first one was a chromosomal intergenic region between an HTH-type transcriptional regulator *CdhR* and the coding region for the Phosphoserine phosphatase 1. This *k*-mer incremented on average 0.63 the logODDS for resistance probability according to SHAP (Table 2; Figure 4). This region was present mainly in isolates from the clonal complex CC258. This is the clonal complex with the highest prevalence of resistance to amikacin in our dataset (73%), which may indicate a different regulation pattern in the isolates of this clade (Figure S1). The second region where the *k*-mer was also present was the coding region for the *ABC-F type ribosomal protection protein Msr(E),* which was associated with amikacin resistance in 90% of cases (n = 38). In addition, the second *k-mer* that contributed the most with an average increment in logODDS of 0.40 was found in the *16S rRNA (guanine(1405)-N(7))-methyltransferase gene* sequence (Table 2; Figure 4) and predicted amikacin resistance in 81% of cases (n = 48).

**Figure 4.**
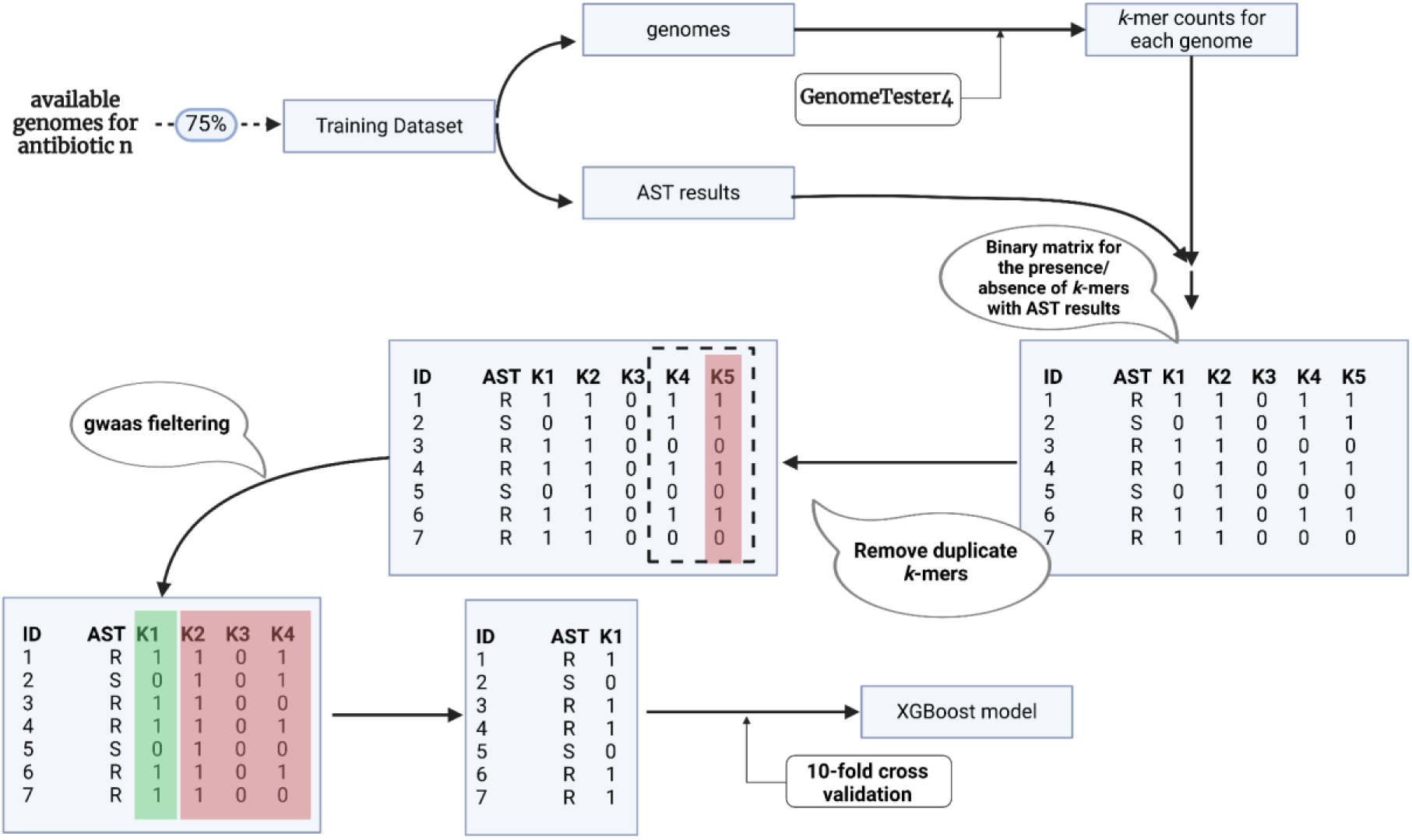
Pipeline used to train the XGBoost models for predicting the resistant phenotype to fosfomycin, amikacin and piperacillin/tazobactam.

For fosfomycin, ResFinder resulted in a high false positive rate (1.00) due to considering *fosA* genes as predictive of resistance. The ML model’s top *k*-mers with high SHAP values were located in genes other than *fosA* (Figure 3). The first *k*-mer was located in an intergenic region of the chromosome, 100 bp downstream the second *k*-mer, which was placed in an hypothetical protein identical to the NCBI protein WP_002887322.1. This region was found in the same contig as the *fosA* gene in 387 genomes, and the distance between the two features ranged between 13 and 23 Kpb, being 13 Kpb in 264 cases. The absence of these *k*-mers was associated with fosfomycin resistance in 43% of cases (n = 478), resulting in an average increment of the logODDS of 0.70 and 0.60 for the first and the second *k-* mer, respectively (Table 2; Figure 4). Additionally, the sixth most important *k*-mer was included in the coding region of a putative tyrosine-protein kinase protein located in the capsular (cps) region that was associated with susceptibility to fosfomycin in 88% of cases (n = 58), leading to an average reduction of the logODDS for the probability of resistance of 0.33 (Table 2; Figure 4).

For piperacillin/tazobactam resistance prediction, the rule-based strategy showed 585 misclassifications due to false positives. Of these, 445 (76%) were produced by the identification of a *bla*_OXA-1_ gene. The most important *k*-mer in the ML model produced an average increment of the logODDS for resistance probability of 0.29 (Table 2; Figure 4). The *k*-mer was annotated within the coding sequence of a *LysR* transcriptional factor, and was associated with piperacillin/tazobactam resistance in 80% of cases (n = 217). Remarkably, this hit was found in genomes from many STs within a contig predicted to be plasmidic. Additionally, the sixth most important *k*-mer was annotated in the gene *dmsB*, responsible for the B chain of the anaerobic dimethyl sulfoxide reductase, and its absence was associated with resistance to piperacillin/tazobactam in 55% of cases (n = 861), with an average increment of the logODDS of 0.14 (Table 2; Figure 4).

## Discussion

In this study we have applied machine learning models to improve the prediction accuracy of AMR phenotypes in *K. pneumoniae* from genome sequences. We have shown that these models are substantially more efficient than rule-based methods, particularly for antibiotics for which the latter have shown limitations. By leveraging on 5,907 *K. pneumoniae* genomes from a diverse dataset that includes both publicly available genomes and a genomic surveillance collection from Spanish hospitals, we have evaluated ML’s predictive power. Our analysis not only provides more accurate predictions but also offers valuable biological insights into the genetic features associated with resistance to three antibiotics, amikacin, fosfomycin, and piperacillin/tazobactam.

The results show a better predictive performance of ML models compared to rule-based strategies for several critical antibiotics. While ResFinder performed well for carbapenems, which are critical last-resort treatments (26), it showed limited effectiveness in predicting resistance to other commonly used antibiotics, especially amikacin, fosfomycin, and piperacillin/tazobactam, which are important first-line treatments (27). This decreased accuracy for predicting amikacin resistance and piperacillin/tazobactam resistance using rule-based methods is consistent with other studies. Ruppé *et al.* reported accuracies of 0.66 and 0.53 for amikacin and piperacillin/tazobactam resistance prediction using ResFinder (6), and Mahfouz *et al*., also using ResFinder, obtained accuracies of 0.40 and 0.58 for these antibiotics (13).

In this study, we explored the genetic determinants that led to inaccurate ResFinder predictions for these antibiotics. For amikacin, most misclassifications in the rule-based strategy were false positives attributed to the detection of the *AAC(6’)-Ib-cr* gene. Removing this gene as a resistance predictor significantly improved the prediction accuracy. The *AAC(6’)-Ib-cr* gene is a variant of the *AAC(6’)-Ib* gene. While *AAC(6’)-Ib* encodes an enzyme that confers resistance to amikacin, the enzyme encoded by *AAC(6’)-Ib-cr* shows reduced affinity for the antibiotic. In *Escherichia coli*, this affinity reduction results in a lower MIC, which falls below the EUCAST breakpoints (*28, 29*). This finding, coupled with the high prevalence of the *AAC(6’)-Ib-cr* gene in *K. pneumoniae,* underscores the need for detailed studies to determine the MIC of this variant in clinical strains.

In the case of fosfomycin, all the genomes were classified as resistant due to the presence of the *fosA* gene, a known determinant of fosfomycin resistance (30). However, while *K. pneumoniae* genomes commonly harbor the *fosA* gene in the chromosome, its expression is insufficient to confer clinical resistance to fosfomycin, explaining the misclassifications (31). Therefore, it is important to incorporate this consideration into rule-based prediction tools when predicting AMR in this pathogen.

For piperacillin/tazobactam, ResFinder predicted resistance with the presence of the *bla_OXA-1_* gene. Although his gene encodes for the beta-lactamase OXA-1, which confers resistance to piperacillin/tazobactam (32, 33), our data showed that *bla*_OXA-1_ presence did not consistently produce clinical resistance, resulting in a high rate of false positives. Similarly to the *AAC(6’)-Ib-cr* variant, further investigations are necessary to determine the MIC of the *bla_OXA-1_* variant in clinical strains.

For these three antibiotics we trained ML models that significantly overperformed the rule-based strategy for predicting the resistant phenotype. These findings are in-line with other studies. Nguyen *et al.* used *k-*mers with the XGBoost algorithm to predict the MIC of different antibiotics in *K. pneumoniae* and found a ±1 tier accuracies of 0.97 for amikacin and 0.78 for piperacillin-tazobactam (15). Similarly, Byeonggyu *et al.* applied convolutional neural networks and obtained raw accuracies of 0.84 for amikacin and 0.68 for piperacillin-tazobactam (34). However, there is a lack of studies benchmarking the prediction of resistance to fosfomycin in *K. pneumoniae* using either ML or rule-based strategies.

In addition to improving prediction accuracy, ML can generate new knowledge about the mechanisms underlying antimicrobial resistance (21). In this study, we used feature importance and SHAP values to extract the most important *k*-mers from each model and extract its functional annotation to explore its implication in resistance mechanisms. For amikacin’s ML model, the *k-*mer that contributed the most to the phenotype was found in the coding region of a *16S rRNA (guanine(1405)-N(7))-methyltransferase* which is known to produce resistance to aminoglycosides (35). Additionally, the most important *k-*mer was present in the *Msr(E)* gene which encodes for a ribosomal protector described as a macrolide resistance producer (36). This gene was not associated with amikacin resistance in the ResFinder database (37). Further studies should detail the role of these genes in *K. pneumoanie* AMR.

The fosfomycin resistance ML model was also interrogated. Among the most important *k*-mers we identified a putative tyrosine kinase involved in the regulation of the capsular polysaccharide synthesis. In addition, we identified a small hypothetical protein whose absence was strongly related to fosfomycin resistance according to SHAP values. These findings provide a hint for investigating additional genetic and molecular determinants of resistance to this antibiotic.

Finally, from the ML model predicting piperacillin/tazobactam resistance we retrieved the anaerobic dimethyl sulfoxide reductase as an important feature determining the phenotype. However, SHAP values for the absence of this feature were bimodal, indicating a possible interaction with other genetic elements. This feature has been found in similar studies as associated with the resistance phenotype to ß-lactams (38). We also identified the most important *k-*mer as a *lysR* transcriptional regulator within an integron with other four genes. Among these genes we identified *glnQ,* which regulates glutamine levels. In *Streptococcus pneumoniae* it has been seen that alterations of glutamine levels can affect susceptibility to some ß-lactams (39). The role of glutamine in *K. pneumoniae* should be addressed.

Our study has several limitations. The computational cost of the analysis restricted our investigation to subsets of data and smaller *k*-mers, which may limit the biological insights that could be derived from larger datasets and longer *k*-mers. Additionally, we had to adopt a categorical classification of resistant (R) and susceptible (S) as the predicted variables due to the variability in AST and MIC results across different clinical settings and years, which may not be directly comparable. Using MIC values instead would have provided more detailed and informative predictions. Moreover, the diversity and representativeness of the dataset might limit the generalization of our findings, because some clonal groups and geographic regions may be overrepresented in both datasets.

In summary, our findings highlight the necessity for a continued exploration of ML applications in the genomic surveillance and prediction of AMR. These models have the potential to significantly improve the prediction of AMR phenotypes in *K. pneumoniae*, particularly in cases where traditional methods are insufficient. Furthermore, they can help us to better understand the genetic mechanisms underlying resistance phenotypes. Continuous evaluation and enhancement of ML techniques will be crucial for effectively integrating them into the clinical practice and public health strategies, including surveillance.

## Methods

### Genomes and AST

We included 1,605 *K. pneumoniae* genome assemblies from clinical isolates collected during the Surveillance of *Klebsiella pneumoniae* in the Comunidad Valenciana (SKPCV) project (40). This project was conducted in 8 different hospitals of the region between January 2017 and February 2020 and was focused on non-susceptible strains to third-generation cephalosporins and/or carbapenems. Additionally, we collected 4,487 available genome assemblies of *K. pneumoniae* in the PATRIC database that had Minimum Inhibitory Concentration (MIC) data (41). Accession numbers of the whole genome collection are provided in Table S3.

Assemblies were characterized using Kleborate (11). To ensure the genome data quality we kept the genomes with a strong species match classified as “strong” by Kleborate with *K. pneumoniae*, a genome size between 5 and 6 Mb and that were assembled in less than 800 contigs.

MICs from the SKPCV were evaluated using VITEK2 whereas those uploaded to PATRIC were mainly obtained from broth microdilution. The antibiotics analyzed included amikacin, ampicillin, aztreonam, cefepime, ceftazidime, ciprofloxacin, fosfomycin, gentamicin, imipenem, meropenem, piperacillin/tazobactam, and tobramycin. MIC values for each antibiotic were classified as susceptible (S) or resistant (R) using the AMR v2.1.0 library for R (42) and the EUCAST 2018 clinical breakpoints (29). Intermediate MICs according to these rules were considered as susceptible.

### Rule based prediction

We used a popular rule-based prediction tool, ResFinder (9), that predicts resistance profiles (resistant/susceptible) based on the identification of antimicrobial resistance genes and point mutations collected in their database. The Resfinder database version used was v2.2.1 and the Pointfinder database version was v4.0.1.

### ML-prediction models

Antibiotics for which the rule-based prediction yielded accuracies below 75% were selected for ML model training. Due to computational constraints, we limited the sample size per antibiotic to a maximum of 1,500 randomly selected samples. Each dataset was then divided into training (75%) and testing (25%) subsets.

To train the models, we used *k*-mer counts (k = 15) extracted from each genome and the AMR phenotype profile (resistant/susceptible). *k*-mers were obtained using GenomeTester4 (43). *k-*mer counts were summarized into a binary matrix where the rows represented the genomes and the columns were the *k-*mers. The matrix was deduplicated, keeping only one k-mer for each unique presence/absence profile. Finally, we filtered the matrix to keep only those k-mers with a possible association with an AMR phenotype (p-value < 0.1). Associations were evaluated using *Fisher’s exact test* corrected with Holm’s method implemented in the stats R library (44). We used the XGBoost R library (45), which implements the Extreme-Gradient Boosting algorithm, to train the models for prediction tasks using decision trees. For each antibiotic, the optimal number of boosting iterations was obtained with a 10-fold cross-validation (Figure 4).

For hyperparameter optimization, we used gentamicin data, due to the number of isolates with AST results (4,015) and balanced proportion of resistant and susceptible isolates (47% and 53%, respectively). We generated 10 random subsets of 500 samples each, which were then divided into training (400 samples) and testing (100 samples) sets. For each subset, we trained models using each combination of learning rate (0.04, 0.05, 0.06 and 0.07) and subsampling percentage (0.3, 0.4, 0.5, 0.6, 0.7, 0.8, 0.9 and 1). This process was replicated 10 times for each sample subset and parameter combination. The selection of the best combination of parameters was based on the Area under the Curve (AUC) distribution for replicates estimated using the R library pROC (46). Means were compared using the Kruskal-Wallis’ test and variances using the Levene’s test, both implemented in the stats R library (44). To limit the complexity of the models and prevent overfitting, we set the maximum tree depth to 4 in all models used.

### ML-based AMR prediction

ML models were used to predict AMR in the SKPCV dataset and PATRIC data. For each model the *k*-mers were fetched using GenomeTester4 (43). Then, an input matrix was calculated for the testing data using the stats R library (44) and predictions were made on the resistant phenotype for the corresponding antibiotic using the XGBoost R library (45). Additionally, optimal probability thresholds for classification were determined using the R library pROC (46).

### Comparison of rule based and ML prediction strategies

Each prediction was marginally analyzed and characterized by computing the true positive rate (TPR), true negative rate (TNR), accuracy, balanced accuracy (bAcc) and F1-score using the R library caret (47), which also computed 95% confidence intervals for accuracies.

To assess if ML models significantly improved the prediction accuracy with respect to the rule-based strategy, we generated 1,000 random pseudoreplicate subsets of 100 samples from the testing data. Then, we estimated the bAcc given the ML prediction and the ResFinder prediction. To determine if the mean difference between the two bAcc distributions was significantly different from zero we used a Wilcoxon’s test implemented in the stats R library (44), taking into account paired data and using a 95% confidence level.

### K-mer interpretation and functional annotation

We analyzed the functional annotation of the most significant *k*-mers identified by the ML predictive models for each antibiotic. To identify the ten *k*-mers with larger impact on phenotype, we relied on the *k-*mer’s importance that depicts how valuable each feature was in the construction of the boosted decision trees. Then, for those *k*-mers we estimated the Shapley Additive Explanation (SHAP) (48) using the R library XGBoost (45). SHAP is a game theoretic approach to compute marginal contributions of each feature to a specific prediction. Additionally, we conducted a detailed functional annotation of the identified *k*-mers. The genome collection was annotated with Prokka (49). Then, we used blastn-short (15) to identify where the selected *k*-mers were located in the genome. We kept only blast matches with a 100% of coverage and identity threshold. The contigs in which the *k*-mers were identified were classified as plasmid or chromosome using mlplasmids (50). Finally, we investigated the *k*-mers that were filtered out in the training process due to duplication in order to identify possible missed features.

The phylogenetic trees in Figures S1 and S2 were built using a core genome at 90% alignment constructed using panaroo (51) and generated using IQ-TREE 2 (52) with a GTR+F+I+G4 model. The trees were rendered using iTOL (53).

## Data availability

The assemblies used in this study are available on NCBI under the biosamples provided in the supplementary table S3. MIC data for the studied antibiotics is also available in the supplementary table S3. The trained models and an easy-to-use implementation to predict new genomes are available in GitHub (https://github.com/SeviJordi/WGS_to_AMR).

## Acknowledgments

This work was funded by projects BFU2017-89594R and PID2021-127010OB-100 from Spanish Ministerio de Ciencia e Innovación, and CIPROM-2021-053 from Generalitat Valenciana.

## References

1. Antimicrobial Resistance Collaborators. 2022. Global burden of bacterial antimicrobial resistance in 2019: a systematic analysis. Lancet 399:629–655.

2. Navon-Venezia S, Kondratyeva K, Carattoli A. 2017. Klebsiella pneumoniae: a major worldwide source and shuttle for antibiotic resistance. FEMS Microbiol Rev 41:252–275.

3. World Health Organization. 2024. WHO bacterial priority pathogens list, 2024: bacterial pathogens of public health importance, to guide research, development, and strategies to prevent and control antimicrobial resistance. World Health Organization.

4. Cai Y, Hoo GSR, Lee W, Tan BH, Yoong J, Teo Y-Y, Graves N, Lye D, Kwa AL. 2022. Estimating the economic cost of carbapenem resistant Enterobacterales healthcare associated infections in Singapore acute-care hospitals. PLOS Glob Public Health 2:e0001311.

5. Boattini M, Bianco G, Charrier L, Comini S, Iannaccone M, Almeida A, Cavallo R, De Rosa FG, Costa C. 2023. Rapid diagnostics and ceftazidime/avibactam for KPC-producing Klebsiella pneumoniae bloodstream infections: impact on mortality and role of combination therapy. Eur J Clin Microbiol Infect Dis 42:431–439.

6. Ruppé E, Cherkaoui A, Charretier Y, Girard M, Schicklin S, Lazarevic V, Schrenzel J. 2020. From genotype to antibiotic susceptibility phenotype in the order Enterobacterales: a clinical perspective. Clin Microbiol Infect 26:643.e1–643.e7.

7. Zankari E, Hasman H, Kaas RS, Seyfarth AM, Agersø Y, Lund O, Larsen MV, Aarestrup FM. 2013. Genotyping using whole-genome sequencing is a realistic alternative to surveillance based on phenotypic antimicrobial susceptibility testing. J Antimicrob Chemother 68:771–777.

8. Stoesser N, Batty EM, Eyre DW, Morgan M, Wyllie DH, Del Ojo Elias C, Johnson JR, Walker AS, Peto TEA, Crook DW. 2013. Predicting antimicrobial susceptibilities for Escherichia coli and Klebsiella pneumoniae isolates using whole genomic sequence data. J Antimicrob Chemother 68:2234–2244.

9. Florensa AF, Kaas RS, Clausen PTLC, Aytan-Aktug D, Aarestrup FM. 2022. ResFinder - an open online resource for identification of antimicrobial resistance genes in next-generation sequencing data and prediction of phenotypes from genotypes. Microb Genom 8.

10. Feldgarden M, Brover V, Gonzalez-Escalona N, Frye JG, Haendiges J, Haft DH, Hoffmann M, Pettengill JB, Prasad AB, Tillman GE, Tyson GH, Klimke W. 2021. AMRFinderPlus and the Reference Gene Catalog facilitate examination of the genomic links among antimicrobial resistance, stress response, and virulence. Sci Rep 11:12728.

11. Lam MMC, Wick RR, Watts SC, Cerdeira LT, Wyres KL, Holt KE. 2021. A genomic surveillance framework and genotyping tool for Klebsiella pneumoniae and its related species complex. Nat Commun 12:4188.

12. Fida M, Cunningham SA, Murphy MP, Bonomo RA, Hujer KM, Hujer AM, Kreiswirth BN, Chia N, Jeraldo PR, Nelson H, Zinsmaster NM, Toraskar N, Chang W, Patel R, Antibacterial Resistance Leadership Group. 2020. Core genome MLST and resistome analysis of Klebsiella pneumoniae using a clinically amenable workflow. Diagn Microbiol Infect Dis 97:114996.

13. Mahfouz N, Ferreira I, Beisken S, von Haeseler A, Posch AE. 2020. Large-scale assessment of antimicrobial resistance marker databases for genetic phenotype prediction: a systematic review. J Antimicrob Chemother 75:3099–3108.

14. Antonopoulos DA, Assaf R, Aziz RK, Brettin T, Bun C, Conrad N, Davis JJ, Dietrich EM, Disz T, Gerdes S, Kenyon RW, Machi D, Mao C, Murphy-Olson DE, Nordberg EK, Olsen GJ, Olson R, Overbeek R, Parrello B, Pusch GD, Santerre J, Shukla M, Stevens RL, VanOeffelen M, Vonstein V, Warren AS, Wattam AR, Xia F, Yoo H. 2019. PATRIC as a unique resource for studying antimicrobial resistance. Brief Bioinform 20:1094–1102.

15. Nguyen M, Brettin T, Long SW, Musser JM, Olsen RJ, Olson R, Shukla M, Stevens RL, Xia F, Yoo H, Davis JJ. 2018. Developing an in silico minimum inhibitory concentration panel test for Klebsiella pneumoniae. Sci Rep 8:421.

16. Nguyen M, Long SW, McDermott PF, Olsen RJ, Olson R, Stevens RL, Tyson GH, Zhao S, Davis JJ. 2019. Using Machine Learning To Predict Antimicrobial MICs and Associated Genomic Features for Nontyphoidal. J Clin Microbiol 57.

17. Kim J, Greenberg DE, Pifer R, Jiang S, Xiao G, Shelburne SA, Koh A, Xie Y, Zhan X. 2020. VAMPr: VAriant Mapping and Prediction of antibiotic resistance via explainable features and machine learning. PLoS Comput Biol 16:e1007511.

18. Lüftinger L, Májek P, Beisken S, Rattei T, Posch AE. 2021. Learning From Limited Data: Towards Best Practice Techniques for Antimicrobial Resistance Prediction From Whole Genome Sequencing Data. Front Cell Infect Microbiol 11:610348.

19. Yang Y, Walker TM, Walker AS, Wilson DJ, Peto TEA, Crook DW, Shamout F, CRyPTIC Consortium, Zhu T, Clifton DA. 2019. DeepAMR for predicting co-occurrent resistance of Mycobacterium tuberculosis. Bioinformatics 35:3240– 3249.

20. Tsang KK, Maguire F, Zubyk HL, Chou S, Edalatmand A, Wright GD, Beiko RG, McArthur AG. 2021. Identifying novel β-lactamase substrate activity through prediction of antimicrobial resistance. Microb Genom 7.

21. Li S, Wu J, Ma N, Liu W, Shao M, Ying N, Zhu L. 2023. Prediction of genome-wide imipenem resistance features in Klebsiella pneumoniae using machine learning. J Med Microbiol 72:001657.

22. Macesic N, Bear Don’t Walk OJ IV, Pe’er I, Tatonetti NP, Peleg AY, Uhlemann A-C. 2020. Predicting Phenotypic Polymyxin Resistance in Klebsiella pneumoniae through Machine Learning Analysis of Genomic Data. mSystems 5.

23. Condorelli C, Nicitra E, Musso N, Bongiorno D, Stefani S, Gambuzza LV, Carchiolo V, Frasca M. 2024. Prediction of antimicrobial resistance of Klebsiella pneumoniae from genomic data through machine learning. PLoS One 19:e0309333.

24. Magiorakos A-P, Srinivasan A, Carey RB, Carmeli Y, Falagas ME, Giske CG, Harbarth S, Hindler JF, Kahlmeter G, Olsson-Liljequist B, Paterson DL, Rice LB, Stelling J, Struelens MJ, Vatopoulos A, Weber JT, Monnet DL. 2012. Multidrug-resistant, extensively drug-resistant and pandrug-resistant bacteria: an international expert proposal for interim standard definitions for acquired resistance. Clin Microbiol Infect 18:268–281.

25. Nguyen M, Brettin T, Long SW, Musser JM, Olsen RJ, Olson R, Shukla M, Stevens RL, Xia F, Yoo H, Davis JJ. 2018. Developing an in silico minimum inhibitory concentration panel test for Klebsiella pneumoniae. Sci Rep 8:421.

26. Livermore DM, Woodford N. 2006. The beta-lactamase threat in Enterobacteriaceae, Pseudomonas and Acinetobacter. Trends Microbiol 14:413–420.

27. Viladomiu AS, Pueyo JM, Suñé EL, Zboromyrska Y, Mondejar PL, López JB. 2022. Guía de terapéutica antimicrobiana: 2022: Mensa Gatell. Antares.

28. Robicsek A, Strahilevitz J, Jacoby GA, Macielag M, Abbanat D, Park CH, Bush K, Hooper DC. 2006. Fluoroquinolone-modifying enzyme: a new adaptation of a common aminoglycoside acetyltransferase. Nat Med 12:83–88.

29. ESCMID-European Society of Clinical Microbiology, Diseases I. eucast: Clinical breakpoints and dosing of antibiotics. https://www.eucast.org/clinical_breakpoints. Retrieved 23 December 2022.

30. Falagas ME, Vouloumanou EK, Samonis G, Vardakas KZ. 2016. Fosfomycin. Clin Microbiol Rev 29:321–347.

31. Kieffer N, Guzmán-Puche J. 2024. The Importance of Genomic Context in Interpreting Fosfomycin Resistance in Klebsiella pneumoniae. Int J Antimicrob Agents 64:107210.

32. Livermore DM, Day M, Cleary P, Hopkins KL, Toleman MA, Wareham DW, Wiuff C, Doumith M, Woodford N. 2019. OXA-1 β-lactamase and non-susceptibility to penicillin/β-lactamase inhibitor combinations among ESBL-producing Escherichia coli. J Antimicrob Chemother 74:326–333.

33. Walkty A, Karlowsky JA, Lagacé-Wiens PRS, Golden AR, Baxter MR, Denisuik AJ, McCracken M, Mulvey MR, Adam HJ, Zhanel GG. 2022. Presence of the narrow-spectrum OXA-1 β-lactamase enzyme is associated with elevated piperacillin/tazobactam MIC values among ESBL-producing Escherichia coli clinical isolates (CANWARD, 2007-18). JAC Antimicrob Resist 4:dlac027.

34. Ryu B, Jeon W, Kim D. 2024. Integrating genomic and molecular data to predict antimicrobial minimum inhibitory concentration in Klebsiella pneumoniae. Sci Rep 14:25951.

35. Wachino J-I, Doi Y, Arakawa Y. 2020. Aminoglycoside Resistance: Updates with a Focus on Acquired 16S Ribosomal RNA Methyltransferases. Infect Dis Clin North Am 34:887–902.

36. Duan Y, Liu S, Gao Y, Zhang P, Mao D, Luo Y. 2022. Macrolides mediate transcriptional activation of the msr(E)-mph(E) operon through histone-like nucleoid-structuring protein (HNS) and cAMP receptor protein (CRP). J Antimicrob Chemother 77:391–399.

37. Zankari E, Hasman H, Cosentino S, Vestergaard M, Rasmussen S, Lund O, Aarestrup FM, Larsen MV. 2012. Identification of acquired antimicrobial resistance genes. J Antimicrob Chemother 67:2640–2644.

38. ValizadehAslani T, Zhao Z, Sokhansanj BA, Rosen GL. 2020. Amino Acid k-mer Feature Extraction for Quantitative Antimicrobial Resistance (AMR) Prediction by Machine Learning and Model Interpretation for Biological Insights. Biology 9.

39. El Khoury JY, Boucher N, Bergeron MG, Leprohon P, Ouellette M. 2017. Penicillin induces alterations in glutamine metabolism in Streptococcus pneumoniae. Sci Rep 7:14587.

40. García-González N, Beamud B, Sevilla-Fortuny J, Sánchez-Hellín V, Vidal I, Rodriguez Díaz JC, Fuster B, Tormo N, Salvador C, Gimeno C, Gomila-Sard B, Giner Almaraz S, Martínez O, Colomina J, Navarro D, Domínguez V, Gonzalez F. 2024. Different Transmission Patterns between Third Generation Cephalosporin and Carbapenem Resistance in Klebsiella Pneumoniae.

41. Wattam AR, Abraham D, Dalay O, Disz TL, Driscoll T, Gabbard JL, Gillespie JJ, Gough R, Hix D, Kenyon R, Machi D, Mao C, Nordberg EK, Olson R, Overbeek R, Pusch GD, Shukla M, Schulman J, Stevens RL, Sullivan DE, Vonstein V, Warren A, Will R, Wilson MJC, Yoo HS, Zhang C, Zhang Y, Sobral BW. 2014. PATRIC, the bacterial bioinformatics database and analysis resource. Nucleic Acids Res 42:D581–91.

42. Berends MS, Luz CF, Friedrich AW, Sinha BNM, Albers CJ, Glasner C. 2022. AMR: An R Package for Working with Antimicrobial Resistance Data. Journal of Statistical Software 10.18637/jss.v104.i03.

43. Kaplinski L, Lepamets M, Remm M. 2015. GenomeTester4: a toolkit for performing basic set operations - union, intersection and complement on k-mer lists. Gigascience 4:58.

44. Team RC. 2022. R: A Language and Environment for Statistical Computing.

45. Tianqi Chen and Tong He and Michael Benesty and Vadim Khotilovich and Yuan Tang and Hyunsu Cho and Kailong Chen and Rory Mitchell and Ignacio Cano and Tianyi Zhou and Mu Li and Junyuan Xie and Min Lin and Yifeng Geng and Yutian Li and Jiaming Yuan. 2024. xgboost: Extreme Gradient Boosting (1.7.7.1).

46. Robin X, Turck N, Hainard A, Tiberti N, Lisacek F, Sanchez J-C, Müller M. 2011. pROC: an open-source package for R and S+ to analyze and compare ROC curves. BMC Bioinformatics 12:77.

47. Kuhn M. 2008. Building Predictive Models in R Using the caret Package. J Stat Softw 28:1–26.

48. Lundberg SM, Erion G, Chen H, DeGrave A, Prutkin JM, Nair B, Katz R, Himmelfarb J, Bansal N, Lee S-I. 2020. From Local Explanations to Global Understanding with Explainable AI for Trees. Nat Mach Intell 2:56–67.

49. Seemann T. 2014. Prokka: rapid prokaryotic genome annotation. Bioinformatics 30:2068–2069.

50. Arredondo-Alonso S, Rogers MRC, Braat JC, Verschuuren TD, Top J, Corander J, Willems RJL, Schürch AC. 2018. mlplasmids: a user-friendly tool to predict plasmid- and chromosome-derived sequences for single species. Microb Genom 4.

51. Tonkin-Hill G, MacAlasdair N, Ruis C, Weimann A, Horesh G, Lees JA, Gladstone RA, Lo S, Beaudoin C, Floto RA, Frost SDW, Corander J, Bentley SD, Parkhill J. 2020. Producing polished prokaryotic pangenomes with the Panaroo pipeline. Genome Biology 21:1–21.

52. Minh BQ, Schmidt HA, Chernomor O, Schrempf D, Woodhams MD, von Haeseler A, Lanfear R. 2020. IQ-TREE 2: New Models and Efficient Methods for Phylogenetic Inference in the Genomic Era. Mol Biol Evol 37:1530–1534.

53. Letunic I, Bork P. 2024. Interactive Tree of Life (iTOL) v6: recent updates to the phylogenetic tree display and annotation tool. Nucleic Acids Res 52:W78–W82.

